# Comparative genomics reveals the emergence of an outbreak-associated *Cryptosporidium parvum* population in Europe and its spread to the USA

**DOI:** 10.1101/2023.09.19.558430

**Authors:** Greta Bellinzona, Tiago Nardi, Michele Castelli, Gherard Batisti Biffignandi, Martha Betson, Yannick Blanchard, Ioana Bujila, Rachel Chalmers, Rebecca Davidson, Nicoletta D’Avino, Tuulia Enbom, Jacinto Gomes, Gregory Karadjian, Christian Klotz, Emma Östlund, Judith Plutzer, Ruska Rimhanen-Finne, Guy Robinson, Anna Rosa Sannella, Jacek Sroka, Christen Rune Stensvold, Karin Troell, Paolo Vatta, Barbora Zalewska, Claudio Bandi, Davide Sassera, Simone M. Cacciò

**Affiliations:** Department of Biology and Biotechnology, University of Pavia, Italy; Department of Comparative Biomedical Sciences, School of Veterinary Medicine, University of Surrey, Guildford, UK; Viral Genetics and Biosecurity Unit (GVB), French Agency for Food, Environmental and Occupational Health Safety (ANSES), Ploufragan, France; Department of Microbiology, Public Health Agency of Sweden, Solna, Sweden; Cryptosporidium Reference Unit, Swansea, UK; Norwegian Veterinary Institute, Ås, Norway; Istituto Zooprofilattico Sperimentale dell’Umbria e delle Marche, Perugia, Italy; Animal Health Diagnostic Unit, Finnish Food Authority, Kuopio, Finland; National Institute for Agricultural and Veterinary Research, Lisbon, Portugal; Ecole Nationale Vétérinaire d’Alfort, Laboratoire de Santé Animale, Maisons-Alfort, France; Department of Infectious Diseases, Unit for Mycotic and Parasitic Agents and Mycobacteria, Robert Koch Institute, Berlin, Germany; National Veterinary Institute, Uppsala, Sweden; National Institute for Public Education, Budapest, Hungary; Finnish Institute for Health and Welfare, Helsinki, Finland; Department of Infectious Diseases, Istituto Superiore di Sanità, Rome, Italy; Department of Parasitology and Invasive Diseases, National Veterinary Research Institute, Pulawy, Poland; Statens Serum Institut, Copenhagen, Denmark; Veterinary Research Institute, Department of Food and Feed Safety, Brno, Czech Republic; Department of Biosciences, University of Milan, Milan, Italy; IRCCS Fondazione Policlinico San Matteo, Pavia, Italy

## Abstract

The zoonotic parasite *Cryptosporidium parvum* is a global cause of gastrointestinal disease in humans and ruminants. Sequence analysis of the highly polymorphic *gp60* gene enabled the classification of *C. parvum* isolates into multiple groups (e.g. IIa, IIc, Id) and a large number of subtypes. In Europe, subtype IIaA15G2R1 is largely predominant and has been associated with many water-and food-borne outbreaks. In this study, we generated new whole genome sequence (WGS) data from 123 human-and ruminant-derived isolates collected in 13 European countries and included other available WGS data from Europe, Egypt, China and the USA (n=72) in the largest comparative genomics study to date. We applied rigorous filters to exclude mixed infections and analysed a dataset from 141 isolates from the zoonotic groups IIa (n=119) and IId (n=22). Based on 28,047 high quality, biallelic genomic SNPs, we identified three distinct and strongly supported populations: isolates from China (IId) and Egypt (IIa and IId) formed population 1, a minority of European isolates (IIa and IId) formed population 2, while the majority of European (IIa, including all IIaA15G2R1 isolates) and all isolates from the USA (IIa) clustered in population 3. Based on analyses of the population structure, population genetics and recombination, we show that population 3 has recently emerged and expanded throughout Europe to then, possibly from the UK, reach the USA where it also expanded. In addition, genetic exchanges between different populations led to the formation of mosaic genomes. The reason(s) for the successful spread of population 3 remained elusive, although genes under selective pressure uniquely in this population were identified.

## Introduction

The genus *Cryptosporidium* (phylum Apicomplexa) currently comprises 46 species and more than 120 genotypes of uncertain taxonomic status (Innes et al. 2020; U. M. Ryan et al. 2021; Tůmová et al. 2023). Although the parasite has a global distribution, cryptosporidiosis represents a high-burden disease in children living in low-income countries, where it is a leading cause of moderate-to-severe diarrhoea (Kotloff et al. 2013), and is associated with long-term negative impacts on childhood growth and well-being (Khalil et al. 2018).

Most *Cryptosporidium* species and genotypes have a narrow host range, suggesting coevolution with their hosts (U. Ryan et al. 2021). Indeed, calibrated phylogenies indicate that much of *Cryptosporidium* diversity originated in the Cretaceous, as was the case for most of the mammals (Garcia-R and Hayman 2016). The mechanisms underlying host adaptation in *Cryptosporidium* are still poorly understood. Several species are known to infect different hosts, including *C. parvum*, *C. felis, C. canis, C. cuniculus, C. ubiquitum, C. meleagridis* and others (Rachel M. Chalmers et al. 2018; Zahedi and Ryan 2020).

With no effective drugs and no vaccine, control of cryptosporidiosis is heavily dependent on the prevention of infection, which has to be informed by a detailed understanding of the epidemiology, population structure, and transmission dynamics of these parasites (Bhalchandra, Cardenas, and Ward 2018; Chavez and White 2018). The epidemiology of human cryptosporidiosis is complex, with transmission occurring indirectly via contaminated food or water, or directly via contact with infected animals or individuals (McKerr et al. 2018). Most human cases are due to *C. hominis*, which is anthroponotic, or *C. parvum*, which is zoonotic (Feng, Ryan, and Xiao 2018). Animal reservoirs, in particular young ruminants, have an essential role in the spillover and spillback of *C. parvum* to humans (Guo et al. 2021).

The most commonly used method for genotyping *C. parvum* isolates is by sequence analysis of the hypervariable gene coding for a 60 kDa glycoprotein 60 (*gp60*), which allowed delineating multiple groups, with IIa, IIc and IId being the most common (Feng, Ryan, and Xiao 2018). In Europe, many IIa subtypes have been identified in humans, and many also circulate among animals. However, a few subtypes appear to predominate, particularly subtype IIaA15G2R1, which is also the most common subtype globally (R. M. Chalmers and Cacciò 2016). The reasons for this high prevalence are unknown.

Recent studies based on whole genome sequence (WGS) comparisons have started to explore the evolutionary genetics of *C. parvum* (Corsi et al. 2023; T. Wang et al. 2022; Feng et al. 2017). In the work of Corsi et al. (2023), analysis of 32 WGSs indicated a clear separation between European and non-European (Egypt and China) isolates, and highlighted the occurrence of recombination events between these populations. Another work analysed 101 WGSs and hypothesised the existence of two ancestral populations, represented by IId isolates from China and IIa isolates from Europe. The authors proposed that the IId and IIa populations recently became sympatric in Europe, and generated hybrid genomes through recombination, possibly influencing biological traits such as host preference (T. Wang et al. 2022).

In this study, we generated WGSs for 123 human-or ruminant-derived *C. parvum* isolates collected across Europe, and retrieved publicly available WGS data of 71 isolates from Europe, Egypt, China and the USA (Corsi et al. 2023; T. Wang et al. 2022; Feng et al. 2017; Hadfield et al. 2015; Troell et al. 2016). Based on the largest comparative study to date, our main aim was to understand the evolution of this important zoonotic pathogen in Europe and in the USA.

## Material and Methods

### 1. Parasite isolates

Table S1 lists the information available for the 195 *C. parvum* isolates from humans and ruminants included in this study. The dataset comprised 123 isolates sequenced in the present study, 71 isolates from previous studies (Hadfield et al. 2015; Troell et al. 2016; Feng et al. 2017; Corsi et al. 2023; T. Wang et al. 2022), and the recently assembled IOWA-ATCC genome (Baptista et al. 2022), which was used as a reference genome.

### 2. Oocyst purification, DNA processing and sequencing

An aliquot of the 123 faecal isolates was used to extract genomic DNA and to identify the species and the *gp60* subtype, using previously published protocols (U. Ryan et al. 2003; Alves et al. 2003). The procedures for DNA purification and extraction are detailed in Corsi et al. (Corsi et al. 2023). In short, oocyst were purified from faecal specimens by immunomagnetic separation, treated with bleach and used for genomic DNA extraction. Genomic DNA was subjected to whole genome amplification (WGA) using the REPLI-g Midi-Kit (Qiagen), according to the manufacturer’s instructions.

For Next Generation Sequencing experiments, about 1 μg of purified WGA product per sample was used to generate Illumina Nextera XT 2 x 150 bp paired-end libraries, which were sequenced on an Illumina NovaSeq 6000 SP platform. Library preparation and sequencing were performed at the ICM (Institut du Cerveau) in Paris, France.

### 3. Data filtering and SNP calling

Raw reads of the 194 isolates were quality-checked and then pre-processed to remove low-quality bases and adapter sequences using Trimmomatic v.0.36 (Bolger, Lohse, and Usadel 2014), with default parameters. A series of sequential steps were then applied to select isolates and SNPs according to multiple criteria (Supplementary Figure 1, Supplementary Table S1).

The presence of *Cryptosporidium* spp. sequences was verified using MetaPhlan v. 3.0.13 (Beghini et al. 2021) and phyloFlash v. 3.4 (Gruber-Vodicka, Seah, and Pruesse 2020). Only isolates showing the presence of *Cryptosporidium* spp. were retained for further analyses.

The *C. parvum* IOWA-ATCC (Baptista et al. 2022) was used as a reference genome to map the filtered reads of each sample with bowtie2 v.2.5.0 (Langmead and Salzberg 2012) with default settings. PCR duplicates were then marked using Picard MarkDuplicates v. 2.25.4 (https://broadinstitute.github.io/picard/). Variant calling (SNPs and indels) was performed using the GATK’s HaplotypeCaller v. 4.2.2.0 (DePristo et al. 2011; Van der Auwera and O’Connor 2020) with default parameters and option -ERC GVCF. SNPs were removed if quality depth <2.0, Fisher strand >60.0, mapping quality <30.0, mapping quality rank-sum test <−12.5, read position rank-sum test <−8.0, and strand odds ratio >3.0.

Read depth and number of missing sites were calculated for each isolates using VCFtools (Danecek et al. 2011), and isolates with a mean read depth <20X were discarded. The GVCFs were then imported into a GATK GenomicDB using the function GenomicsDBImport, and a combined VCF was created using the GATK GenotypeGVCFs function.

To maximise the quality, SNPs were further filtered using bcftools based on the following criteria: biallelic SNPs, quality score >30, allele depth >20, minor allele frequency >0.005, and missing ratio <0.5.

The moimix R package (https://github.com/bahlolab/moimix) was then used to estimate multiplicity of infection. The FWS statistic, a type of fixation index to assess the within-host genetic differentiation, was calculated on the filtered SNPs. In pure isolates with haploid genomes, FWS is expected to approach unity. Isolates with FWS< 0.95 were excluded, as they were likely to represent multiple infections (Manske et al. 2012). Examples of infections with estimated multiplicity of infection =1 or >1 are presented in Supplementary Figure 2.

Cleaned mapped reads were assembled using Unicycler v.0.5 (Wick et al. 2017) with the --linear_seqs 8 option, which accounts for the presence of eight linear chromosomes in the reference assembly. Isolates with a genome size <8 Mb (the size of the reference genome is 9.1 Mb) were discarded, thus leading to the final dataset (Supplementary Table 1).

SNP differences were calculated using snp-dists v0.8.2 (https://github.com/tseemann/snp-dists) and visualised using the R package heatmap.2. The number of SNPs in non-overlapping windows of 1 kb across each chromosome was counted using vcftools (--SNPdensity) (Danecek et al. 2011), and visualised using the R package ggplot2 (Wickham 2016).

To compare the SNP density between each chromosome we used the pairwise comparisons for proportions test implemented in R, and the probability (p) values were adjusted using the Bonferroni correction.

### 4. Phylogenetic and population structure analyses

To ensure proper rooting of the tree inferred from genomic SNPs, we first generated a tree based on orthologous genes from the 141 *C. parvum* isolates of the final curated dataset, and used *Cryptosporidium hominis* TU502 (GCA_001593465.1) as outgroup. The gene sequences of the *C. hominis* isolate and of the reference genome *C. parvum* IOWA-ATCC were downloaded from CryptoDB (Puiu et al. 2004). The AUGUSTUS algorithm (Stanke et al. 2006) was locally trained on the *C. parvum* IOWA-ATCC genome and then used to predict coding sequences for all the remaining 140 isolates. A set of 195 genes, which have been used previously for phylogenomic analyses of Apicomplexa (Mathur, Wakeman, and Keeling 2021), was searched using BLASTp on the orthogroups identified by OrthoFinder v2.5.4 (Emms and Kelly 2019) in our dataset. Of these, orthologs of 179 genes were identified. Each ortholog was aligned with Muscle 5.1 (Edgar 2004) and concatenated. A maximum likelihood (ML) tree was inferred on the concatenated alignment according to the model indicated by modeltest-ng (HKY+F+I, BIC criteria) (Darriba et al. 2020) with RAxML v.8.2.12 (Stamatakis 2014), with 100 bootstrap pseudo-replicates.

Next, a concatenated set of *C. parvum* genomic SNPs was created by converting the VCF into a FASTA file (https://github.com/edgardomortiz/vcf2phylip). A ML phylogenetic tree was inferred with RAxML v.8.2.12 (Stamatakis 2014) using the GTR+G model, as indicated by modeltest-ng (Darriba et al. 2020), with ascertainment bias correction and 100 bootstrap pseudo-replicates. The same procedure was applied separately on the SNPs located into each of the eight chromosomes, to obtain individual chromosome phylogenies.

Population structure analysis was performed with ADMIXTURE v1.3.0 (Pritchard, Stephens, and Donnelly 2000), with the number of populations tested (K) ranging from 1 to 12. Phylogenetic networks were generated by using the Neighbor-Net algorithm implemented in SplitsTree v.5 (Huson and Bryant 2006).

Pairwise Identity By Descent (IBD) was calculated using a hidden Markov model (Schaffner et al. 2018), and relatedness networks were generated using the R package igraph (Csardi et al., 2006).

### 5. Recombination analyses

The sequence of each chromosome was reconstructed for each isolate by editing the reference IOWA-ATCC sequences according to the corresponding filtered SNPs using the GATK’s FastaAlternateReferenceMaker function (Van der Auwera and O’Connor 2020). Then, multiple sequence alignments of each chromosome were analysed by the Recombination Detection Program software, version 5 (RDP5) (Murrell et al., 2015) using five algorithms (RDP, Geneconv, Bootscan, MaxChi, and Chimæra) implemented in this software. Only events supported by at least three algorithms and with a p-value cut-off of 10e-5 were considered significant.

### 6. Population Genetic Analyses

Tajima’s D values were calculated using snpR (Hemstrom and Jones 2023) in non-overlapping windows of 10 Kb across each entire chromosome. Pairwise divergence (Dxy) and intra-population nucleotide diversity (π) were calculated in genomic windows of 50 kbp sliding by 25 kbp (https://github.com/simonhmartin/genomics_general). The Fixation index (Fst) for each population was computed in windows of 1 kb (https://github.com/simonhmartin/genomics_general).

Decay in Linkage Disequilibrium (LD) was estimated with PopLDdecay 3.42 (Zhang et al. 2019), measuring *r*^2^ between SNPs until 300 kb. The values were computed comparing the mean values of 100 pseudoreplicates, each one composed by 10 isolates extracted randomly.

### 7. Selective pressure analyses

The phylogenetic tree inferred from genomic SNPs was labelled according to the population structure using the dedicated tool of Hyphy v.2.5.50 (Murrell et al. 2015). Then, we determined whether a gene was subjected to positive selection using Hyphy with the BUSTED algorithm on the respective gene sequences from the reconstructed chromosomes (see above). Genes with a p-value < 0.05 were considered statistically significant.

### 8. Comparison of putative virulence genes

A set of 55 putative virulence genes (Dumaine et al. 2021) was retrieved. These genes include members of small gene families characterised by possessing specific protein domains (MEDLE, WYLE, GGC, FLGN, SKSR and mucins) and by having N-terminal signal peptides. The corresponding protein sequences were identified in the assembly of each isolate using BLAST. The results were manually curated, and multiple protein alignments were generated. The presence and distribution pattern of amino acid substitutions were investigated manually.

## Results

### Quality Control and Sample Selection

We started from an initial collection of WGS from 194 *C. parvum* isolates (including 123 newly sequenced isolates and 72 isolates retrieved from public databases). In order to get a robust foundation for reliable inferences, we performed a careful selection based on multiple criteria, including the level of contamination from non-target organisms, mean read depth, multiplicity of infection, and genome assembly quality (more details are provided in the Materials and Methods section and in Supplementary Table S1). This stringent selection process yielded a final dataset of 141 isolates (including 88 newly sequenced isolates and 52 publicly available isolates), and the reference IOWA-ATCC, which were used for downstream analyses.

Importantly, the final dataset comprised isolates from four continents (Africa, Asia, Europe, and North America) and from the two major zoonotic *gp60* groups (IIa and IId). Detailed information regarding the dataset composition can be found in the Supplementary Table S1.

### Genetic variability among isolates

By comparing the 140 isolates to the IOWA-ATCC reference, a total of 45,663 single nucleotide polymorphisms (SNPs) and 18,909 insertion/deletions (InDels), were identified. We filtered the SNPs in order to include only high-quality, biallelic SNPs (see Methods), which reduced the number to 28,047. The SNPs distribution was non-random at the level of individual chromosomes, with statistically significant higher SNP density observed at chromosomes 1 and 6 (p-value < e-05) (Supplementary Table S2), and with an enrichment in subtelomeric regions compared with internal regions of the chromosomes (Supplementary Figure 3).

### Phylogenetic analysis and population structure

To provide a preliminary overview of the global evolutionary relationships among *C. parvum* isolates, we inferred phylogeny based on a defined set of 195 orthologous genes, and included *C. hominis* as an outgroup (Supplementary Figure 4). We observed that all isolates from Europe and the USA formed a highly supported, monophyletic clade (hereafter, the “Western” lineage), whereas isolates from China and Egypt appeared to have diverged earlier.

Next, to get a finer description of the relationships among the 141 *C. parvum* isolates, we inferred a ML tree based on the concatenated set of 28,047 biallelic SNPs (Figure 1), using the root determined in the tree based on orthologous genes. The large-scale topology was consistent with the orthologous genes-based phylogeny. Corroborating indications from previous studies (Corsi et al. 2023; T. Wang et al. 2022), we observed that the host species and *gp60* subtypes were “scattered” along the tree, the latter indicative of limited predictive power for the deep phyletic relationships within *C. parvum*. However, a partial correlation with the geographical origin was found, as isolates sampled from the same region/country, from a single farm, or collected within a narrow temporal window, formed evident subclusters (e.g., all but one of the Finnish isolates formed a monophyletic clade). Interestingly, all isolates from the USA formed a fully supported monophyletic clade that was nested within European isolates and showed an intriguing sister group relationship with a clade of isolates from the UK.

**Figure 1.**
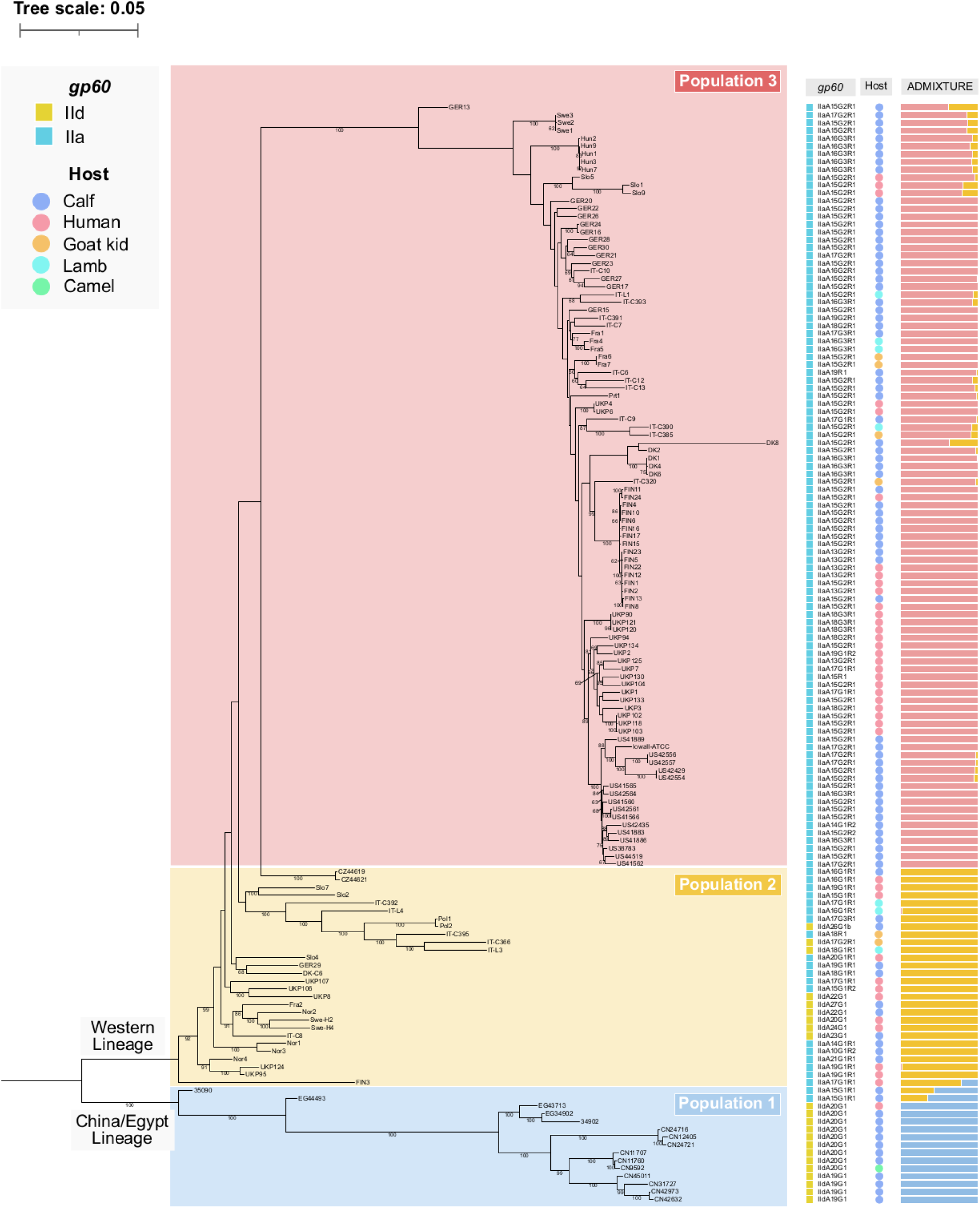
Maximum Likelihood tree based on a set of 28,047 biallelic SNPs. Only bootstrap values >60 are shown. Information about host, *gp60* subtype and results of ADMIXTURE are mapped on the phylogeny.

We then investigated population structure using ADMIXTURE and identified k=3 as the most probable number of populations, in overall agreement with phylogeny (Figure 1). Population 1 encompassed all the Chinese and Egyptian isolates (15), population 2 included a non-monophyletic group of a minority of European isolates (28/109), while population 3 comprised the majority of the European isolates (81/109) and all those from the USA (17), which together formed a monophyletic clade.

While the three populations were genetically very distinct (Figure 1), admixed isolates were also evident, most notably the IIa isolates from Egypt, and in the European isolates FIN3, GER13 and DK8. A phylogenetic network showed several connections between isolates from different populations, suggestive of recombination events (Supplementary Figure 5).

### Recombination analyses

To infer recombination events that may have contributed to the formation of mosaic genomes, we conducted a comprehensive analysis for each chromosome, and performed SNP-based phylogeny, pairwise divergence (Dxy), ADMIXTURE (Zhou, Alexander, and Lange 2011), and SplitsTree (Huson and Bryant 2006). RDP5 (Martin et al. 2015) analyses were performed to provide statistical evidence for putative recombination events.

At chromosome 1, phylogenetic analysis showed the IIa isolates from Egypt (35909 and EG4493) cluster with population 2 and not with population 1, in contrast with the topology based on all genomic SNPs. An inspection of the SNP distribution revealed a mosaic pattern in which the IIa Egyptian isolates are either very similar to the IId isolates from Egypt and China (population 1) or to the IIa and IId isolates from Europe (population 2). Indeed, in the region spanning position 755,934 to 768,672 (about 15 kb), 260 SNPs are found in population 1 (including isolates 35909 and EG4493), while populations 2 and 3 have very limited genetic variability. Immediately after this block, the IIa isolates from Egypt are essentially identical to those from population 2 until position 823,729 (about 55 kb), while the IId isolates differ from the reference genome by 560 SNPs in this region. Among the genes in the latter region, several encode for proteins with signal peptides (e.g., members of the SKSR and CpLSP gene families). This mosaic structure is confirmed by the results of a SplitsTree analysis.

Furthermore, we observed that isolate FIN3, a human-derived isolate from Finland with a history of travel to the Canary Islands, occupied a position between populations 1 and 2 in the phylogenetic analysis, and showed signs of admixture and loops connecting it to population 1. Indeed, in a 50 kb region spanning position 824,800 to 874,170, the isolate FIN3 shared 443 SNPs with the IId isolates from population 1. This region contained several genes encoding for proteins with signal peptides (e.g., members of the SKSR and CpLSP gene families, cgd1_140, cgd1_150, cgd1_160). Therefore, FIN3 is a hybrid that resulted from a recombination event that involved population 1, as further supported by the results of a SplitsTree analysis.

At chromosome 2, in a 210 kb region spanning from position 384,000 to 594,000, the isolate DK8 (calf isolate from Denmark, belonging to population 3) shares 460 SNPs with isolates from population 2 and, less so, population 3. This region contains more than 50 genes, among which the presence of a member of the secreted GGC gene family and of the insulinase-like peptidase family can be noted.

At chromosome 4, a typical mosaic structure is evident in the first 8 kb adjacent to the 5’ telomere. In this region, the IId isolates from China (except those from Hebei and Shanghai), all Egyptian isolates and the European isolates from Norway (calf isolate Nor1), Slovenia (human isolates Slo1, Slo2 and Slo9) and Italy (lamb isolates IT-C392 and IT-L3) shared about 270 SNPs, and differ from all other isolates of populations 2 and 3, which are essentially identical to the reference genome. Four genes are located in this subtelomeric region, all encoding for uncharacterized proteins.

At chromosome 6, in the first 18 kb adjacent to the 5’ telomere, the isolate Ger-13 (calf isolate from Germany, belonging to population 3) shares about 200 SNPs with isolates from populations 1 and 2, while all other population 3 isolates are essentially invariant, being identical to the reference genome. Five genes are located in this subtelomeric region, four encoding for uncharacterized proteins and one for an IMP dehydrogenase/GMP reductase. Therefore, Ger-13 is a hybrid that resulted from a recombination event that involved population 1, as further supported by the results of the SplitsTree analysis.

At chromosome 8, in a 30 kb region spanning position 210,000 to 240,000, the isolate DK8 (calf isolate from Denmark, belonging to population 3) shares 107 SNPs with isolates from population 2 and 3, while the remaining isolates from population 3 are largely invariant. Five genes are located in this region, and encode for two uncharacterized proteins, a protein with putative membrane domain, and a protein with AP2/ERF domain.

### Genomic variability at the population level

We observed that out of the 28,047 SNPs identified in the entire dataset of 140 genomes, only 1,243 (4.4%) were shared by the three populations, while the majority was specific to a single population (Supplementary Figure 6).

We calculated the pairwise SNP distances among the 141 isolates and observed the smallest distances in population 3 (range 3 to 2528 SNPs, average, 892 SNPs). On the other hand, larger SNP distances were observed in population 2 (range 56 to 3532 SNPs, average 1930 SNPs) and in population 1 (range 60 to 4437 SNPs, average 2113 SNPs). Considering inter-population variation, we observed that population 1 exhibited the greatest genetic divergence from both populations 2 and 3, with an average of 5,867 and 5,241 SNPs, respectively, while population 2 and population 3 displayed a lower average SNP distance (3,847 SNPs).

Furthermore, we observed 13 clusters of highly similar genomes (defined as having < 50 SNPs) (Figure 2) all belonging to population 3, encompassing from 2 to 8 isolates. Notably, all clusters were formed by isolates from known outbreaks or from epidemiologically linked cases. As examples, the human-derived isolates UKP102, UKP103 and UKP118 were from an outbreak that occurred in March 2016, while isolates UKP90, UKP120 and UKP121 were from a distinct outbreak that occurred in April 2016. Another cluster of highly similar genomes (i.e., differing for < 20 SNPs) was formed by six Hungarian calf isolates (Hun1, Hun2, Hun3, Hun7 and Hun9), collected at a single farm from the Pest county at multiple but short time intervals (May-June 2020), thus representing clearly epidemiologically linked cases, and a possible outbreak. Another cluster comprised three Swedish calf isolates (Swe1, Swe2, Swe6) collected at a single farm in the same year. In all these cases, a very high genomic similarity was observed (pairwise SNP distance < 20 SNPs), and the respective isolates formed monophyletic, highly supported clusters in the phylogenetic analysis (Figure 1). Although the isolates were not specifically collected to address this question, our data suggest that a threshold of 50 SNPs may be used to identify highly related *C. parvum* strains, which may serve as an appropriate cut-off to confirm suspected outbreaks at the genomic level.

**Figure 2.**
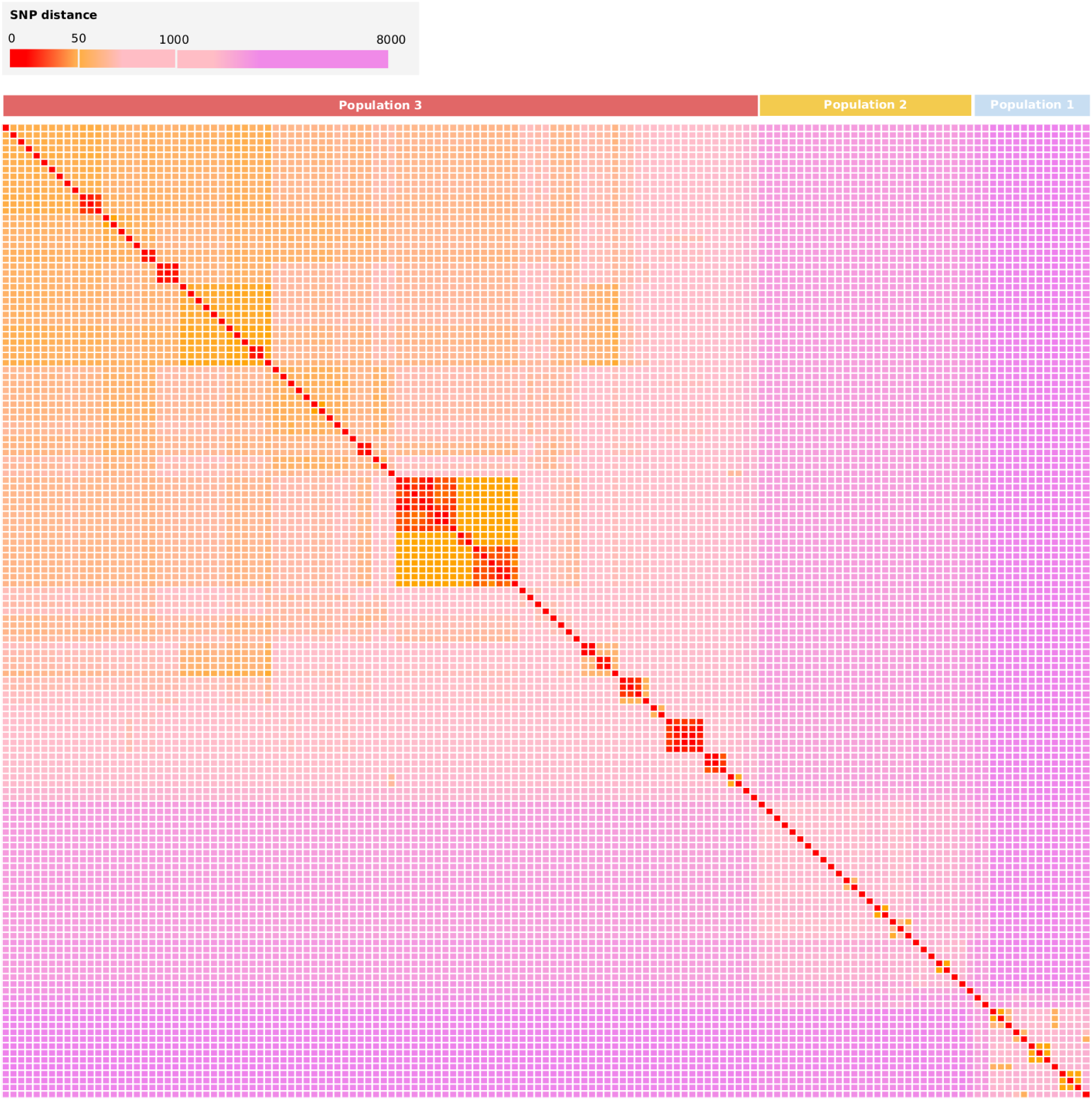
Heatmap illustrating the SNP distance among the 141 isolates analysed. The order of the isolates reflects the position they occupy in the SNP-based phylogenetic tree. The colour code is shown in the legend on the top.

To further investigate relationships among isolates within each population, we undertook an identity by descent (IBD) analysis, and constructed relatedness networks at 90% and 80% (i.e., where the fraction of shared IBD is greater than 90% or 80%). As shown in Supplementary Figure 7, networks were formed by isolates from outbreaks and from single farms, as expected, but also by isolates from specific geographic areas, a result compatible with geographically structured populations. Notably, networks were observed within population 3 (Hungary, Finland, USA/UK, Germany/France) and population 1 (China, Egypt), but not for population 2.

### Within-Population Genetic Indices

To gain further insight into the genetic differentiation of the two populations in the “Western” lineage (i.e., population 2 and 3), we calculated nucleotide diversity (π), linkage disequilibrium (LD) decay, and Tajima’s D.

Assessing nucleotide diversity (π) within each population unveiled notable distinctions. Population 3 exhibited lower diversity compared to population 2 both at the entire genome level (respective means π = 0.039 and π = 0.073) and also when analysing single chromosomes (Figure 3). This reduced genetic variation is consistent with a recent origin of population 3, and can be explained by various factors, such as genetic drift (e.g., population bottlenecks) or selective sweeps.

**Figure 3.**
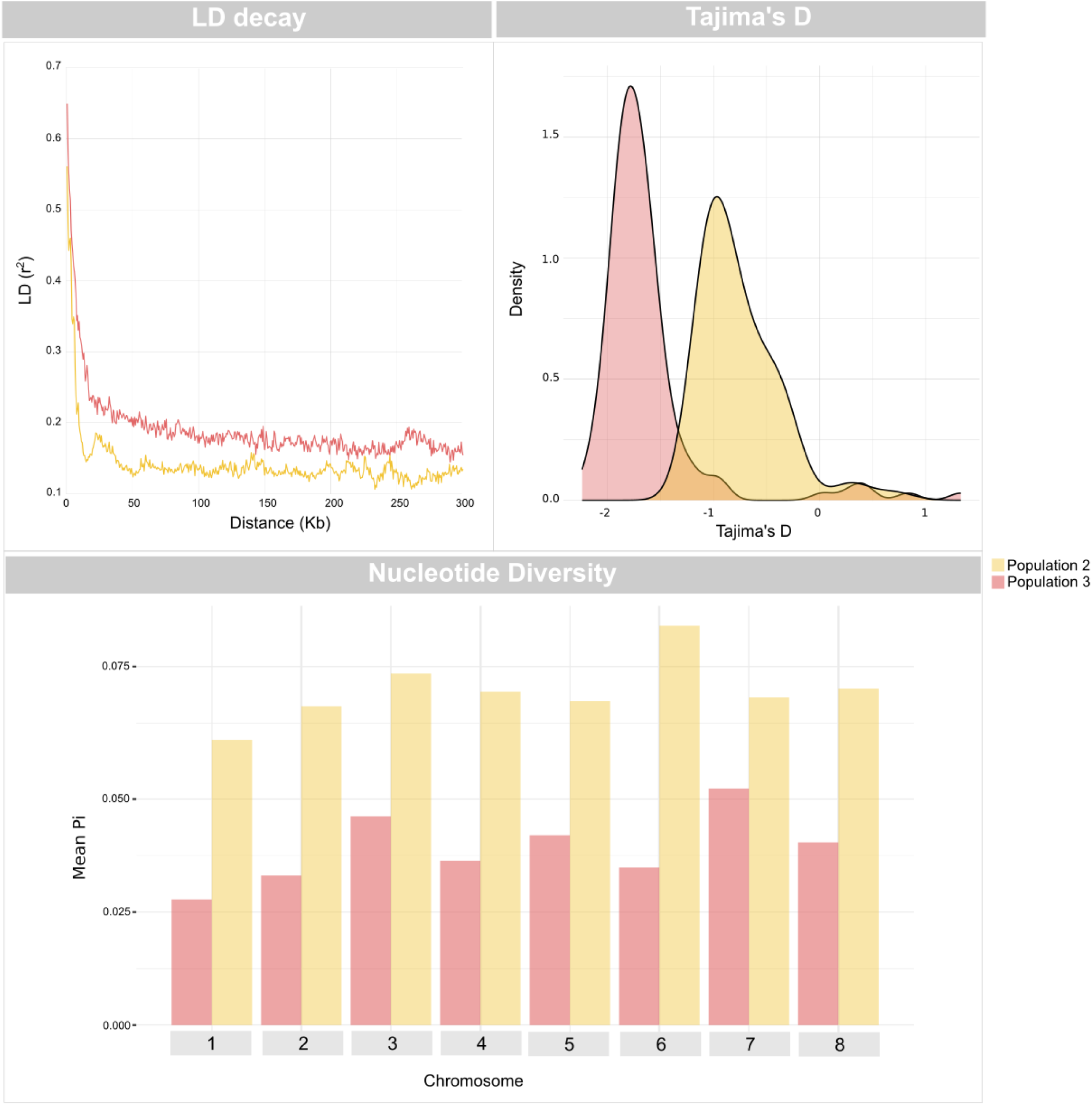
LD decay, distribution of Tajima’s D values and nucleotide diversity (π) in population 2 and population 3. Nucleotide diversity is shown with the mean Pi for each chromosome. Chromosome-specific results are provided in Supplementary Figure 8.

We then examined LD decay and found that population 3 had a slower decay than population 2 (Figure 3), reinforcing the concept of a more recent origin of population 3.

Finally, we observed that the distribution of Tajima’s D values in population 3 is skewed towards negative values (Figure 3), indicating an excess of rare polymorphisms. This skewness was less pronounced in population 2 (Figure 3).

### Genomic differences between population 2 and population 3

We investigated patterns of genetic variation between the two “Western” populations by first screening a set of 55 putative virulence factors involved in the host-parasite interplay (e.g. those encoding for mucin-like glycoproteins, thrombospondin-related adhesive proteins, secreted MEDLE family proteins, insulinase-like proteases and rhomboid-like proteases). We did not find any presence/absence pattern differentiating the two “Western” populations, neither considering amino acid substitutions.

To investigate differences between population 3 and population 2 at the genome-wide level, we calculated the fixation index (Fst) in 1 kb windows (Figure 4). This allowed detection of genomic regions putatively under selection and we inspected the genes present in such regions. By focusing on the top 1% of the total Fst values (i.e., applying a cut-off of 0.91), we identified 79 regions (Figure 4) and highlighted candidate genes under selection according to their function in Figure 4 (Supplementary Table 3).

**Figure 4.**
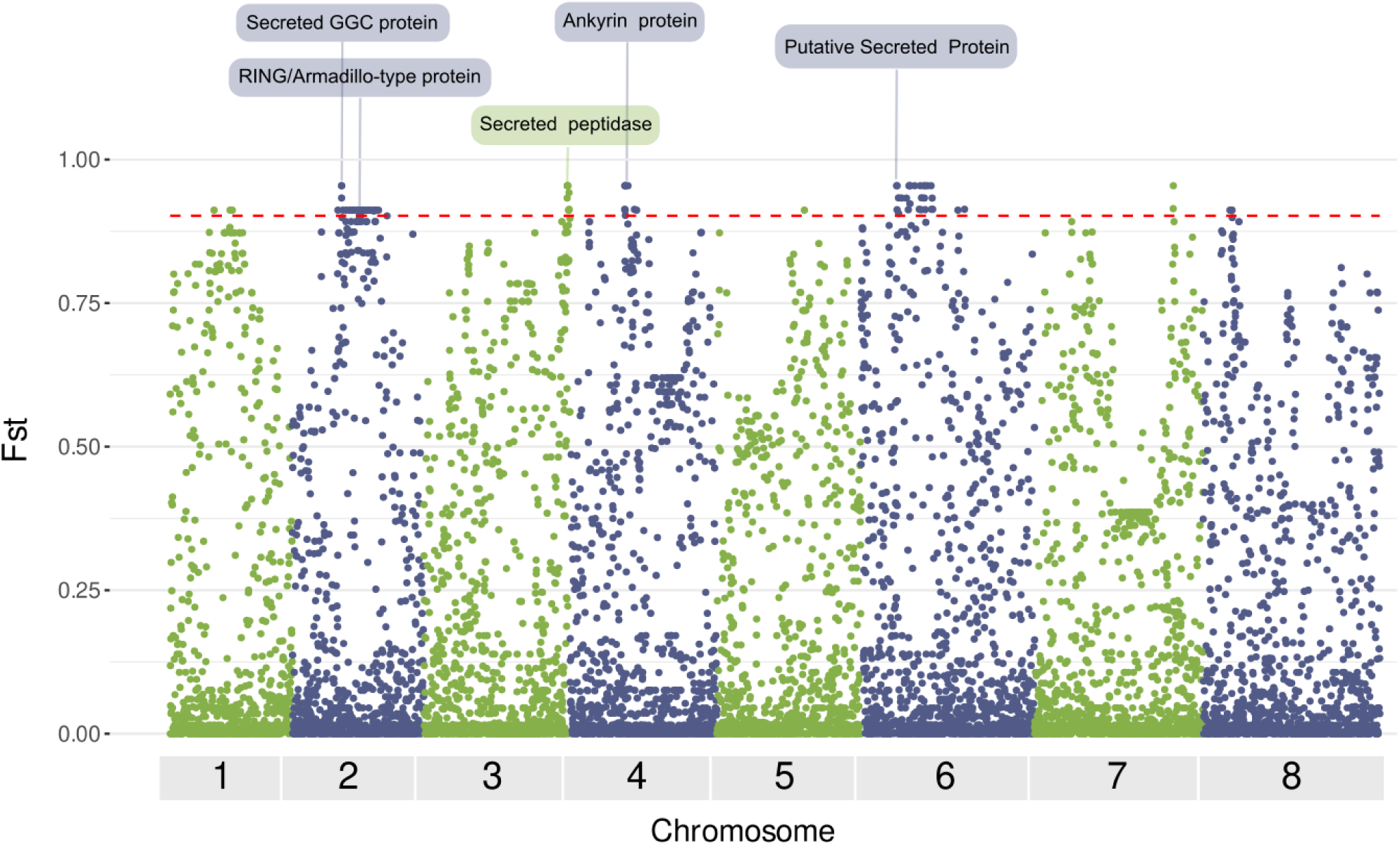
Manhattan plot of genome-wide Wright’s Fst values, calculated in genomic regions of 1 kb, comparing population 2 and population 3. Fst values are shown on the y axis, and genomic positions on the x axis. The dotted red line represents the cut-off value of the top 1%, equal to 0.91. Genes overlapping with the BUSTED analysis and with a function potentially associated with virulence or host-pathogen interaction are highlighted.

Then, we aimed to identify genes in which at least one nucleotide position exhibited significant (p < 0.05) signs of selective pressure. Out of the 3385 annotated genes in the reference IOWA-ATCC, 228 appeared to be under selective pressure in population 3. To test whether these genes were under selective pressure exclusively in population 3, we extended our analysis to population 2. We found that 176 of 228 genes were under selection pressure exclusively in population 3 and not in population 2. Most of these genes (49/176) were annotated as "hypothetical proteins," while a few (8/176) were "putative secreted proteins" (Supplementary Table S3).

Notably, 16 proteins were identified in both the analyses, i.e., Fst statistics on genomic regions and selective pressure in single genes (Supplementary Table S3).

## Discussion

*C. parvum* is the most prevalent zoonotic pathogen within the genus *Cryptosporidium*, and a global cause of diarrheal disease in humans and ruminants (Kotloff et al. 2013). This species is widespread in industrialised countries, including Europe (Cacciò and Chalmers 2016), but also in the Middle East (Hijjawi et al., 2022). Despite the recognised impact on human health and livestock production, no effective drugs or vaccines are available for controlling *C. parvum* infections. An urgent need for new control tools has been repeatedly underlined (Chavez and White 2018; Khan and Witola 2023; Rahman et al. 2022). Recent WGS studies have started to provide insights into the genetics of *C. parvum*, proposing a role of recombination events in the evolution of this species and have allowed to identify a number of genes under positive selection, potentially involved in host-parasite interactions (Corsi et al. 2023; T. Wang et al. 2022).

In this study, we conducted the most extensive comparative genomic analysis of *C. parvum* to date by generating WGS data from human-and ruminant-derived isolates collected in 13 European countries (n=127). Additionally, publicly available WGS data from Europe, Egypt, China, and the USA (n=71) were included (Hadfield et al. 2015; Troell et al. 2016; Feng et al. 2017; Corsi et al. 2023; T. Wang et al. 2022) (Supplementary Table S1). We filtered the initial dataset to obtain a thoroughly cleaned and curated dataset of 141 isolates (including the reference IOWA-ATCC genome), ensuring a robust foundation for reliable genomic analyses. Consistent with earlier findings (Corsi et al. 2023; T. Wang et al. 2022; Baptista et al. 2022), the overall genetic variability was modest, as just 28,047 biallelic high-quality SNPs were identified across the 141 genomes analysed. Phylogenetic analyses using both orthologous genes and SNP data provided robust evidence for the presence of two distinct lineages (Figure 1). One lineage (China/Egypt Lineage) consisted of all Chinese (IId) and Egyptian (IIa and IId) isolates, which was well distinct from the second lineage where all European (IIa and IId) and USA (IIa) isolates are grouped (“Western” lineage; see Fig. 1).

Considering that recombination has been, and still is, a fundamental driver of the genomic evolution of *C. parvum* (Corsi et al. 2023; T. Wang et al. 2022) and other *Cryptosporidium* species (Nader et al. 2019; Huang et al., 2023), we then focused on tracing these events. We described two clear events at chromosomes 1 and 4 (Figures or Table), involving isolates from different populations and hosts, with representatives of population 1 being the putative minor parents (i.e., the donors). Intriguingly, the very same event on chromosome 4 has already reported on a more limited dataset (Corsi et al 2023), while we provided extensive support for a recombination event on chromosome 1 that differed from that reported in Corsi et al. (2023). Additional evidence for the existence of mosaic genomes was obtained from network and admixture analyses of single chromosomes (Supplementary data and Figure). We observed SNP distribution patterns compatible with genetic exchanges, although the precise reconstruction of the events could not be achieved.

A deeper focus on the population structure showed that the “Western” lineage can be divided in two subgroups, namely population 3, a monophyletic group formed by all the USA and most of the European isolates, and population 2, a paraphyletic group that included the remaining European isolates.We propose that population 2 was the ancestral and more heterogeneous European population (including both IIa and IId isolates), and that population 3 (including only IIa isolates) has evolved more recently from it. Our hypothesis is strongly supported by the overall lower genetic diversity (π), negative Tajima’s D values and a slowed decay in LD in population 3 compared to population 2.

Our reconstructions indicate that population 2, which includes both IIa and IId groups, is ancestral in Europe. Considering the relatively limited admixture herein evidenced with extra-European isolates, it seems reasonable to hypothesise that this coexistence of IIa and IId lineages in Europe could date back to ancient introductions of this parasite from the Middle East, which is indeed one of the first areas in which livestock breeding originated (Beja-Pereira et al. 2006; Chessa et al. 2009). This is consistent with previous reconstructions based on the greater diversity of IId subtypes in Asia (R. Wang et al. 2014), and with the coexistence of IIa and IId in Egypt and several other Middle Eastern countries as well (Hijjawi et al. 2022).

A deeper focus on population 3 showed that all the USA isolates (from nine different States) form a monophyletic clade, indicating a single event of introduction from Europe, likely from the UK (Figure 1), and a subsequent expansion in the United States. Historical data (“The Introduction of Cattle into Colonial North America” 1942; Ficek 2019) and studies investigating the ancestry of New World cattles (McTavish et al. 2013; Delsol et al. 2023), suggest that this event should be relatively recent, as most of the import of livestock, particularly cattle, into the Americas occurred from the XVII to the XIX century by Portuguese and Spanish colonists, and during the Victorian Age by Britains (Ficek 2019; McTavish et al. 2013)).

Our results are consistent with *gp60* molecular typing data that identified only IIa subtypes in USA isolates (Jann et al. 2022). A parallel could be drawn with the recent emergence and rapid predominance of the *C. hominis* IfA12G1R5 subtype in the USA (Huang et al., 2023). In this case, however, the emergent lineage originated from successional recombination events involving North American, East African, and European populations (Huang et al., 2023), while in the case of the *C. parvum* population 3, the data supports a single introduction in the country.

Moreover, we found that the clusters of isolates showing high genome similarity (<50 SNPs) all belonged to population 3 (Figure 2), including all those from known outbreaks. Thus, we investigated at genome-wide level the hypothesis of a selective advantage in population 3, which may explain its higher prevalence and association with water-and foodborne outbreaks. While admittedly speculative, the most interesting result comes from a combination of population statistics and phylogeny-based statistical tests on gene sequences, which allowed to identify 16 candidate proteins under positive selection only in population 3. Interestingly, the candidates include genes encoding for secreted proteins, such as ankyrin repeat-containing proteins, which have been shown in *Toxoplasma* to be involved in cell invasion (Long et al. 2017), and a RING/Armadillo-type fold domain containing protein that in *Plasmodium falciparum* mediates the motility of the parasite, essential for fertilisation and transmission (Straschil et al. 2010). While the exact functions of these genes and the biological implications of our observations require further investigation, their identification opens avenues for understanding the mechanisms underlying the selective advantage. Other non-mutually exclusive explanations should be explored, including variation in copy number of genes encoding virulence factors (Xu et al., 2019), their differential expression, or a higher capacity to withstand standard water treatments and persist longer in the environment while maintaining infectivity.

## Conclusions

Our study provides new insights into the epidemiology and evolution of *C. parvum*, with the description of a phylogenetic group that we indicated as the Western lineage. Within this Westen lineage, we observed the presence of two sympatric populations in Europe, and demonstrated that one has recently expanded to become predominant in young ruminants and humans, and then, likely from the UK, reached and spread into the USA. All isolates of the virulent and hyper transmissible IIaA15G2R1 subtype and all outbreak strains belonged to this recently expanding population, suggesting a selective advantage. Investigation of the genes under selective pressure identified a number of candidates with potential roles in the interaction with the host. Overall, our findings allow us to propose a scenario for the evolution of *C. parvum* and to pose focused questions for future research on this parasite.

## Acknowledgements

We would like to thank Dr. Christian Seyboldt (Friedrich Loeffler Institute, Germany) and Dr. Ernst Grossmann (Aulendorf state veterinary diagnostic centre, Germany) for generous provision of original samples. We also would like to thank Elke Radam (Robert Koch Institute, Germany) for excellent technical assistance.

## Author contributions

S.M.C. conceived the study. G.B., T.N., M.C. and G.B.B. performed the bioinformatics analyses. A.R.S. and S.M.C performed the bench work. S.M.C., C.B., G.B., T.N., M.C. and D.S. wrote the manuscript with input from all authors. Authors read and approved the final manuscript.

## Ethics approval and consent to participate

Not applicable

