## Supplementary figures and legends for "Comparative genomics reveals the emergence of an outbreak-associated *Cryptosporidium parvum* population in Europe and its spread to the USA"

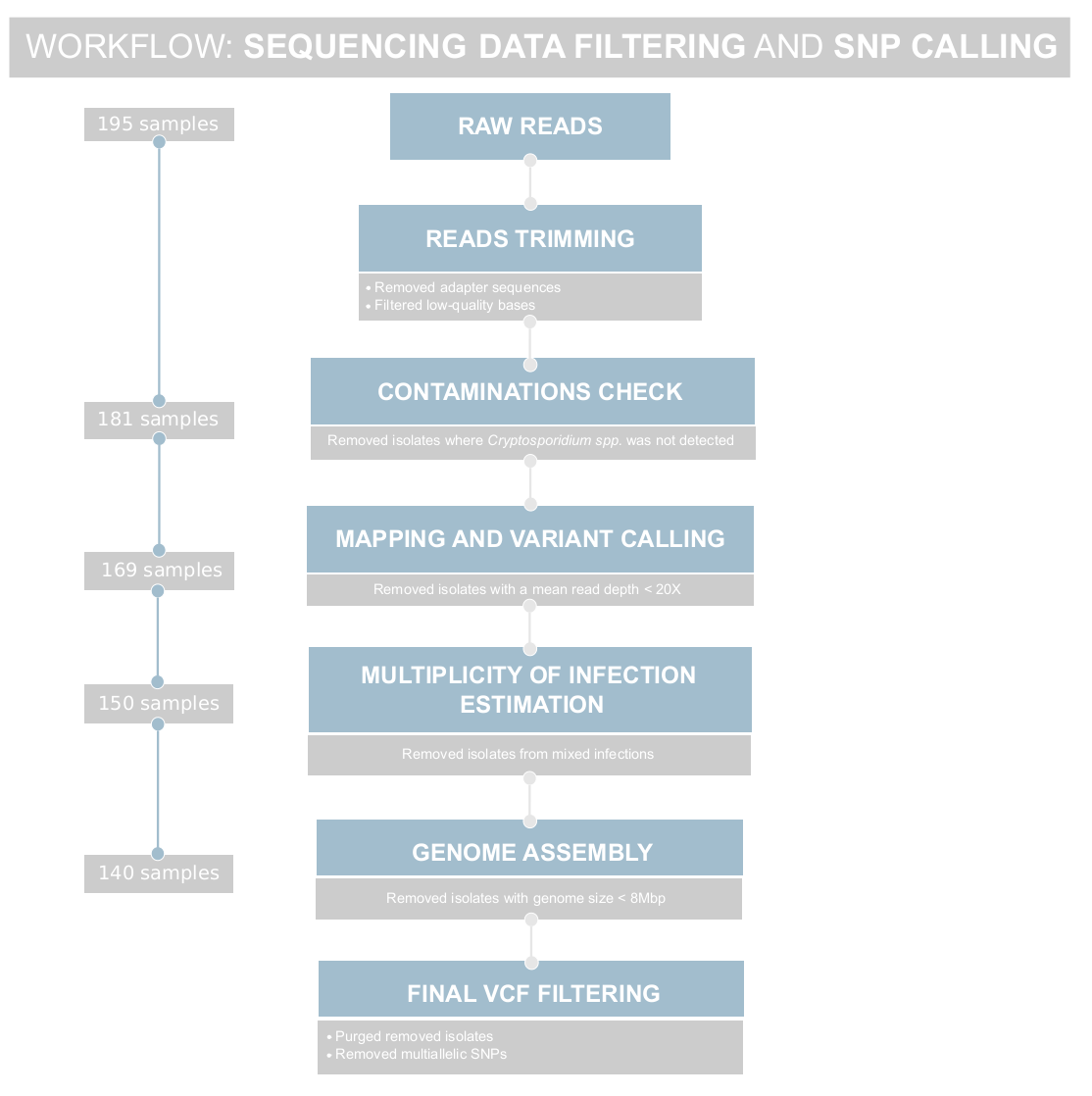
**Figure S1**. **Workflow for data filtering and SNP calling**. The figure outlines the sequential steps employed in data processing to derive the ultimate refined dataset used for the downstream analysis. The left side of the diagram enumerates the count of samples at each filtering step, underscoring the progressive refinement of the dataset. Supplementary Table 1 provides additional details about the specific samples excluded during the various filtering steps.

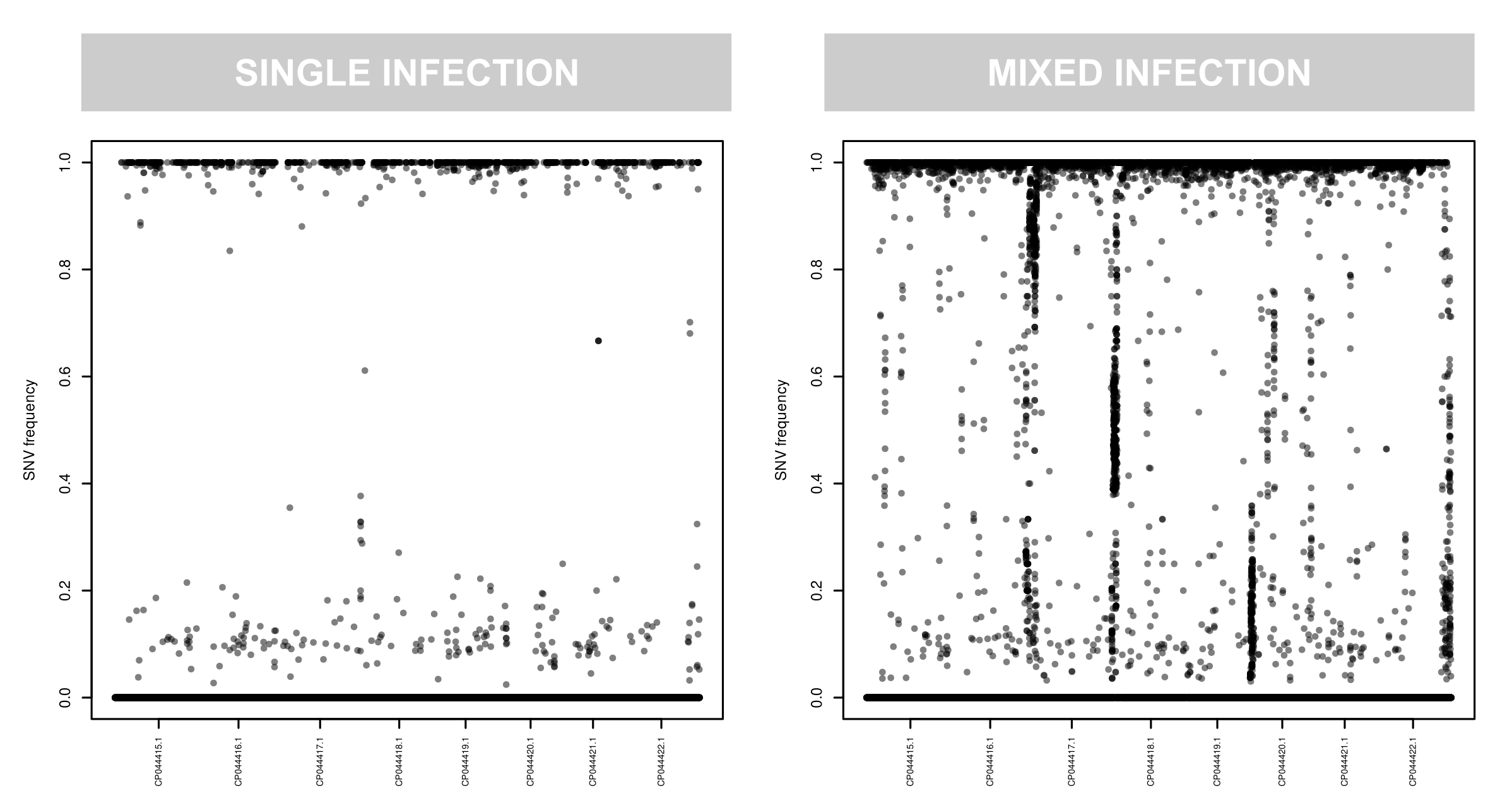
**Figure S2**. **Estimating the presence of mixed infection**. On the left an example isolate that is from a single-clone infection (FIN22). On the right an example for an isolate likely from a multiple-clone infection (EG44409) and that was subsequently excluded from the analysis.

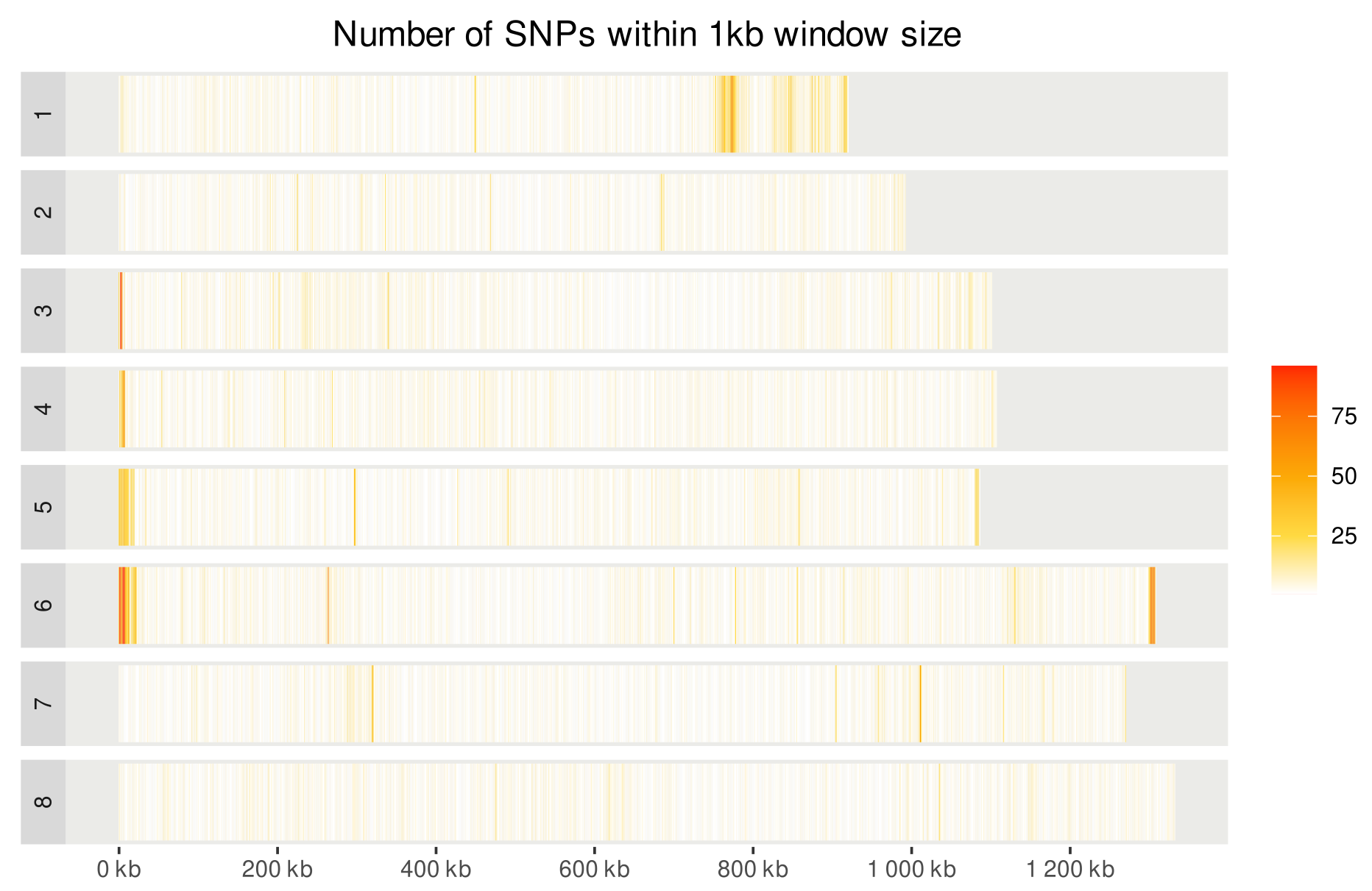

**Figure S3**. **SNP density along the chromosomes**. We counted the number of SNPs in non-overlapping windows of 1 kb across each chromosome. Red correspond to high density SNP regions.

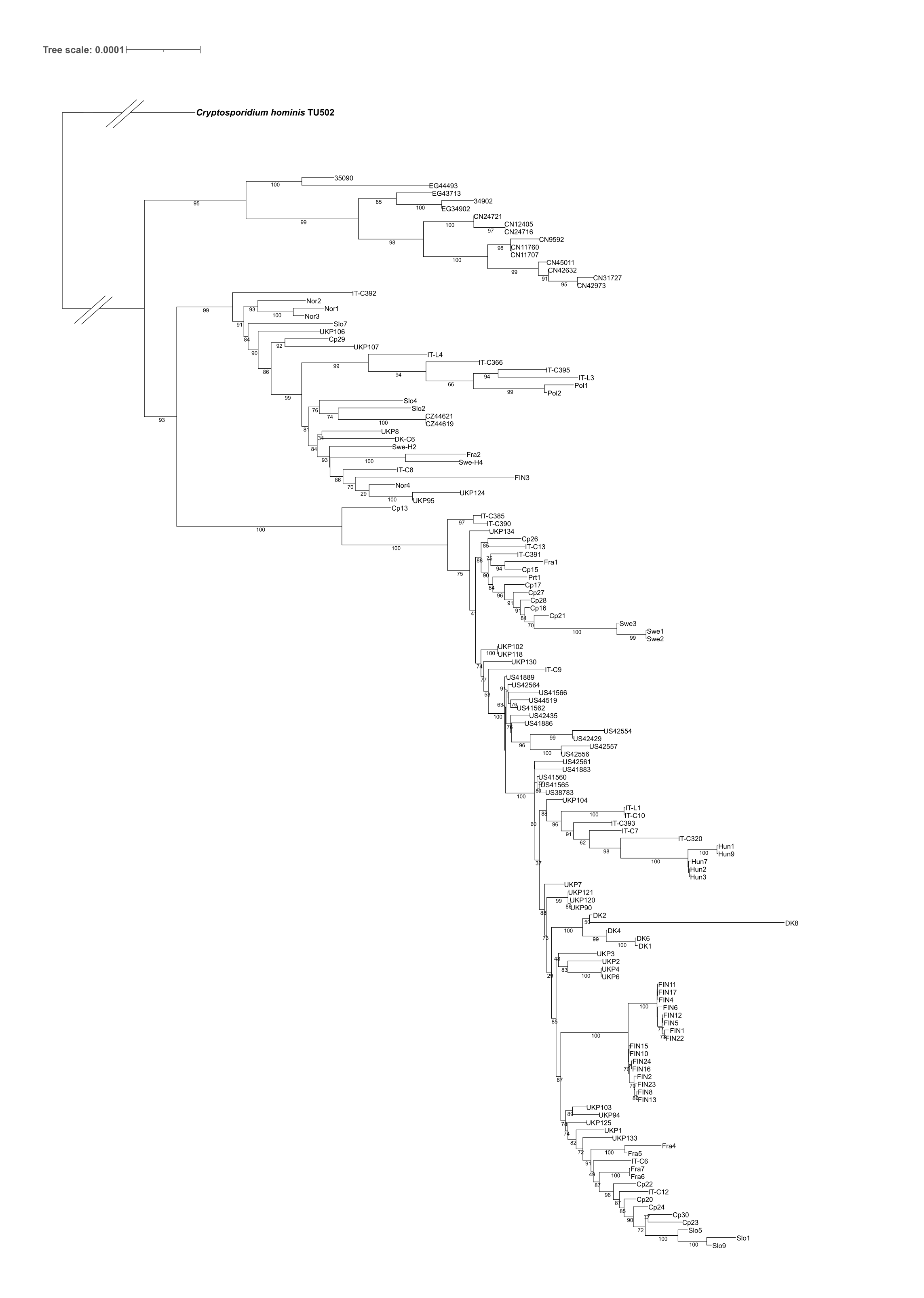

**Figure S4**. Maximum Likelihood (ML) phylogenetic tree inferred on a set of 179 genes. *Cryptosporidium hominis* TU502 was used as outgroup to root. The branch length leading to the outgroup was shortened for presentation purposes.

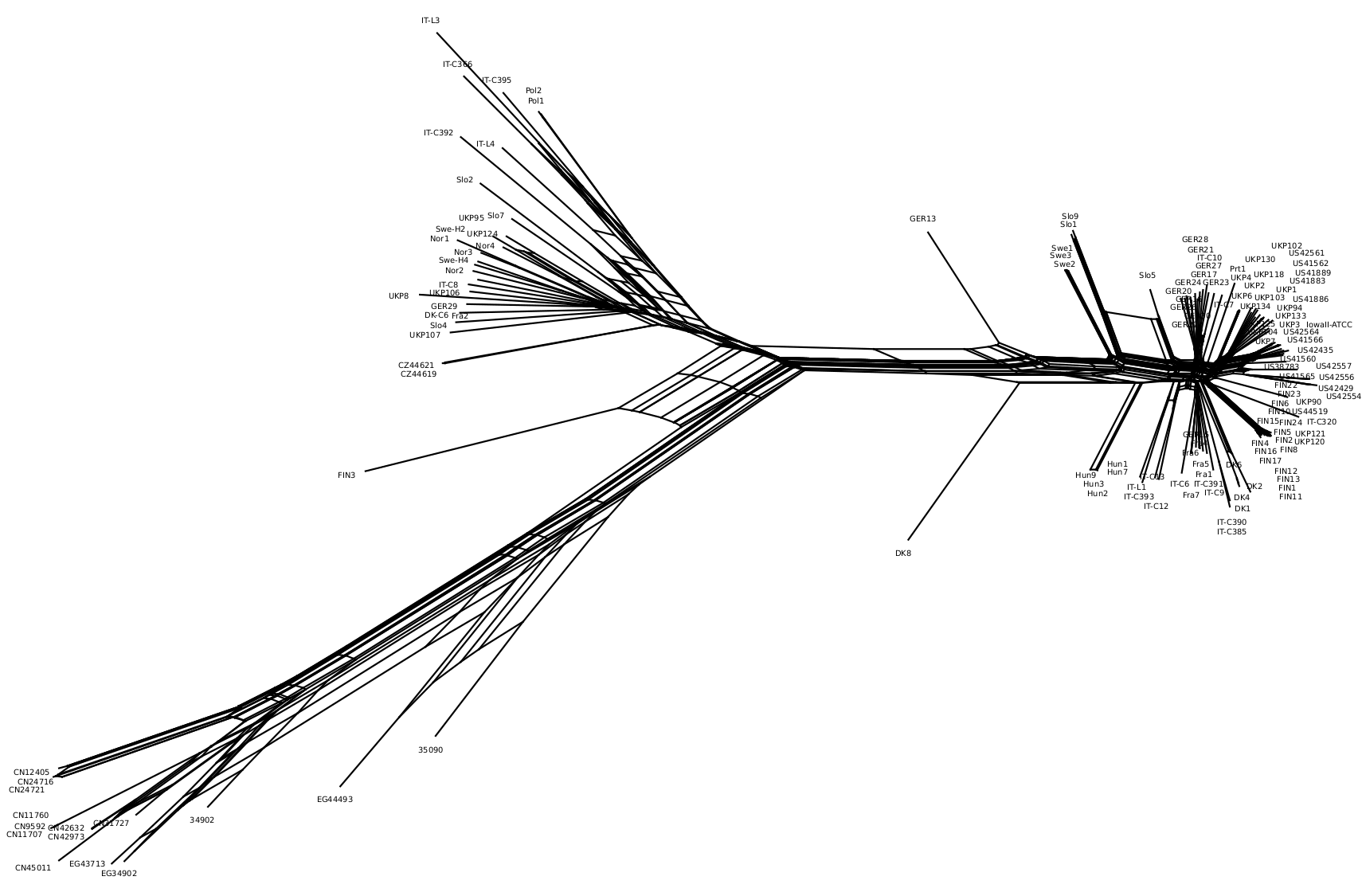
**Figure S5**. Phylogenetic network generated using SplitTree.

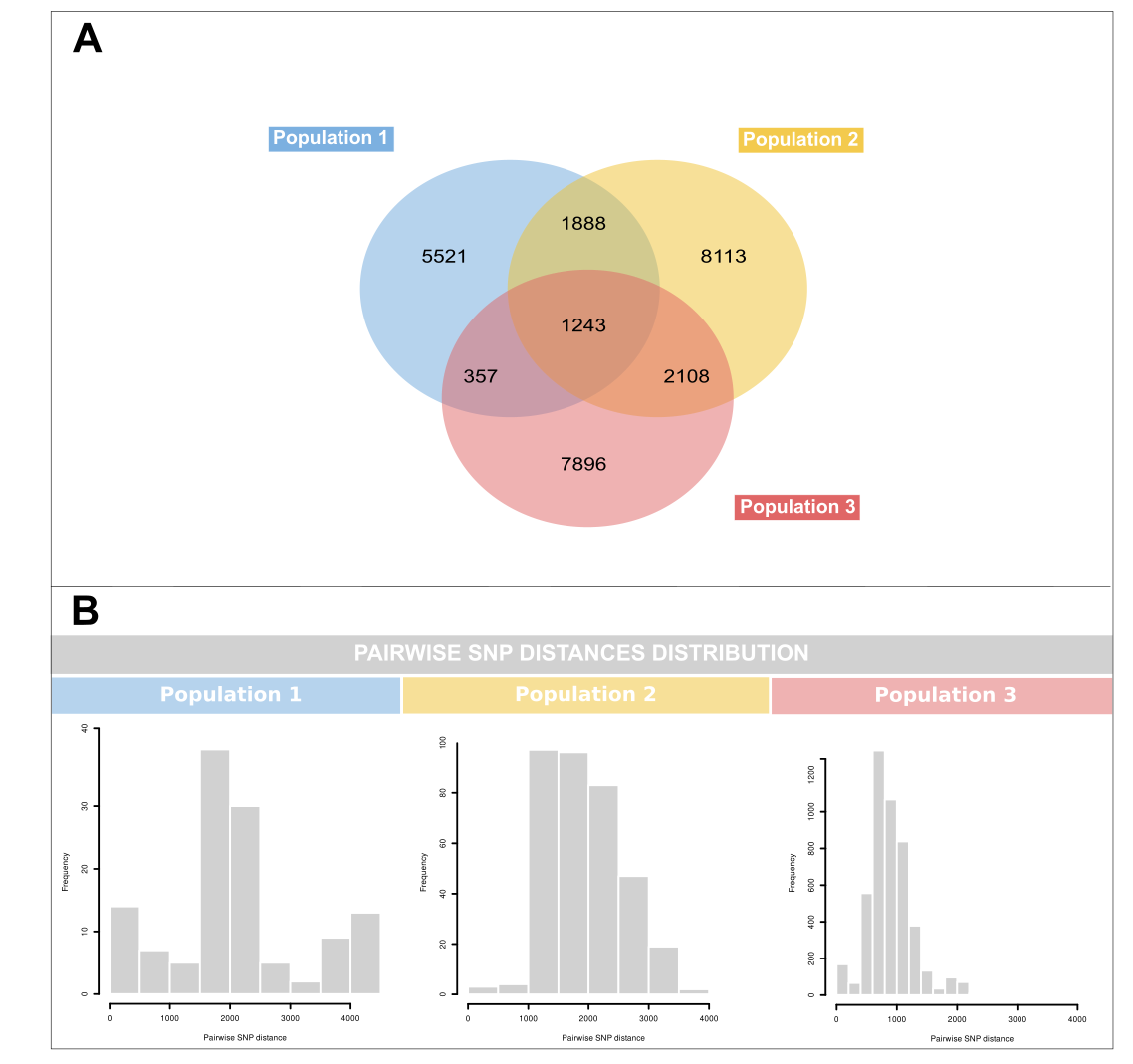

**Figure S6**. **Distribution of shared and population-specific SNPs**. A) Venn Diagram B) Histogram of the pairwise SNP distances distribution within each Population.

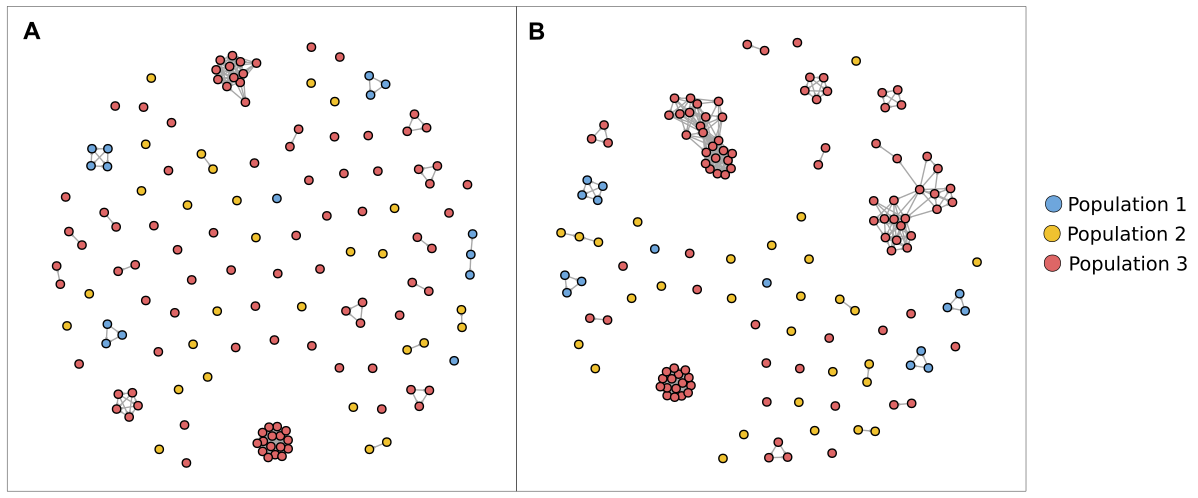
**Figure S7**. **Relatedness network for pairs of isolates identified as having high proportions of IBD sharing**. Each node identifies a unique isolate and an edge is drawn between two isolates if they share more than A) 90% of their genome IBD, and B) 80% of their genome IBD. Individuals are coloured according to the ADMIXTURE population.
